# Sex hormones underlying 17a-Estradiol effects on neuroinflammation

**DOI:** 10.1101/2020.05.26.117689

**Authors:** Lucas K. Debarba, Hashan Jayarathne, Richard A. Miller, Michael Garratt, Marianna Sadagurski

**Author notes:** Corresponding authors: Marianna Sadagurski, Department of Biological Sciences, Integrative Biosciences Center, Wayne State University, 6135 Woodward, Detroit, MI 48202, Michael Garratt, Department of Anatomy, School of Biomedical Sciences, University of Otago, Dunedin 9016, New Zealand.

## Abstract

17-α-estradiol (17aE2) treatment extends lifespan in male mice and can reduce neuroinflammatory responses in the hypothalamus of 12-month-old males. Although 17aE2 improves longevity in males, female mice are unaffected, suggesting a sexually dimorphic pattern of lifespan regulation. We tested whether the sex-specific effects of 17aE2 on neuroinflammatory responses are mediated by sex hormones and whether hypothalamic changes extend to other brain regions in old age. Manipulating sex hormone levels through gonadectomy, we show that sex-specific effects of 17aE2 on age-associated gliosis are brain region-specific and are partially dependent on gonadal hormone production. 17aE2 treatment started at 4 months of age protected 25-month-old males from hypothalamic inflammation. Castration prior to 17aE2 exposure reduced the effect of 17aE2 on hypothalamic astrogliosis. By contrast, sex-specific changes in microgliosis with 17aE2 were not significantly affected by castration in males. While 17aE2 treatment had no effect of hypothalamic astrocytes or microglia in intact females, ovariectomy significantly increased the occurrence of hypothalamic gliosis evaluated in 25-month-old females, which was partially reduced by 17aE2. In the hippocampus, both male and female gonadally-derived hormones influenced the severity of gliosis and the responsiveness to 17aE2 in a region-dependent manner. The male-specific effects of 17aE2 correlate with changes in hypothalamic ERα expression, highlighting a receptor through which 17aE2 could act. The results of this study demonstrate that neuroinflammatory responses to 17aE2 are partially controlled by the presence of sex-specific gonads. Interactions between sex-steroids and neuroinflammation could, therefore, influence late-life health and disease onset, leading to sexual dimorphism in aging.

## Introduction

Aging is characterized by increased neuroinflammatory responses in various brain regions, contributing to age-associated cognitive impairment, memory loss, neurodegenerative diseases, and metabolic imbalance. These neuroinflammatory responses are associated with a marked increase in the number of activated glial cells, specifically the astrocytes and the microglia (Valles *et al.* 2019). Microglia are the resident phagocytes of the innate immune system of the brain that produce inflammatory cytokines in response to stress, disease, or normal aging (Villa *et al.* 2019). During the aging process, activated microglia induce the formation of neurotoxic reactive astrocytes that can contribute to neuronal loss and drive the progression of neurodegeneration (Liddelow *et al.* 2017). With age, there is an alteration in astrocytes phenotype. Astrocytes become more pro-inflammatory and contribute to the inflammatory responses in the aging brain. Due to the astrocytes active interactions and secretory products, aged astrocytes could negatively afflict other CNS cells (Zamanian *et al.* 2012). Our recent studies have demonstrated that age-associated gliosis is reduced in the long-lived mouse models, suggesting that a reduction in neuroinflammatory responses may attenuate the aging process (Sadagurski *et al.* 2015).

Targeting brain immune responses and inflammatory glial cells hold promise for the treatment of neurological diseases and the aging brain. Recent studies have provided evidence that pharmacological interventions that extend mouse life span also reduce age-associated neuroinflammation. Treatment with acarbose (ACA), an anti-diabetes drug, and 17-α-estradiol (17aE2), an optical isomer of 17-β-estradiol (Zhurova *et al.* 2009), causes a consistent extension in male lifespan, but lifespan changes in females are either smaller (acarbose) or do not occur (17aE2). Similarly, we have recently demonstrated that ACA and 17aE2 significantly reduce age-associated neuroinflammatory responses in males but not in females in the hypothalamus, the brain region responsible for metabolic regulation and energy homeostasis. The causes for these sexually dimorphic neuroinflammatory responses to drugs treatment are not clear.

There is substantial sexual dimorphism in the number of glia cells in the brain, occurring during brain development and throughout the adult lifespan. Previous studies have demonstrated that microglial number, morphology, and neuroinflammatory responses are dependent on age and brain region, as well as on hormonal and environmental factors (Grabert *et al.* 2016; Bennett *et al.* 2018). Microglia are likely to be responsive to male and female sex hormones (Kodama & Gan 2019). In particular, it is established that immune functions and inflammatory responses of microglia are significantly mitigated by estrogens (Vegeto *et al.* 2003; Villa *et al.* 2016). Similarly, astrocytes exhibit sex-specific gene expression profiles in different brain regions and are implicated in estrogen-dependent neuroprotection (Acaz-Fonseca *et al.* 2014). Given the emerging evidence that hypothalamic inflammation and microglia activity can influence systemic metabolism and aging, these sex differences in glia abundance and activity could lead to sexual dimorphism in metabolic responses outside of the brain.

Recently, Garratt et al. have demonstrated that male-specific metabolic responses associated with 17aE2 treatment are specifically dependent upon gonadal hormone production (Garratt *et al.* 2017). Male castration before treatment initiation eliminated the insulin and glucose sensitivity induced by drug treatment (Garratt *et al.* 2017). These hormonally-dependent sexspecific effects of 17aE2 for males suggest a potential mechanism of 17aE2 action that requires male-gonadal hormone production. These effects could still involve the estrogen signaling pathway, and possible interaction with the estrogen receptor (ER), but could be linked to steroid metabolism, particularly since 17aE2 leads to a male-specific increase in other estrogen steroids in the liver, particularly sulfated forms of estradiol (Garratt *et al.* 2018). 17aE2 and estradiol both have the capacity to bind to ER, but with lower affinity than 17βE2 (Harrison *et al.* 2014). In support, it has been recently demonstrated that 17aE2 inhibits inflammation via estrogen receptor alpha (ERα) in cultured cells (Santos *et al.* 2017).

Recently, Stout et al. (Stout *et al.* 2016) demonstrated reductions in inflammatory markers in adipose tissue in mice treated with 17aE2, suggesting that 17aE2 may be a useful pharmacological intervention to treat metabolic imbalance and inflammation with aging. Moreover, in a subsequent study, it was shown that some of the short-term metabolic responses to 17aE2, particularly feeding behavior on a high-fat diet, are dependent on the functional presence of pro-opiomelanocortin (POMC) neurons (Steyn *et al.* 2018), a subset of neurons found in the arcuate nucleus of the hypothalamus (ARC). This indicates that hypothalamic actions of 17aE2 are involved in some of the metabolic responses to this drug, at least in the context of obesity. In this study, we evaluated the effect of 17aE2 and sex hormones on neuroinflammatory responses in old 25-month-old mice with a focus on brain regions that are sensitive to age-related neurological changes and metabolic imbalance.

## Results

### Effect of sex hormones on astrogliosis in 17aE2 treated mice

We have previously demonstrated that 17aE2 reduces hypothalamic astrogliosis and microgliosis in a sex-specific manner in male but not female 12-month-old mice. Sex hormones have been linked to sex differences in lifespan (Brooks & Garratt 2017), and can influence glia responsiveness and sex-specific gene expression (Schwarz & Bilbo 2012). To test whether the effects of 17aE2 on age-associated neuroinflammation are dependent on gonadal hormones, we assessed age-associated gliosis in the hypothalamus and in the hippocampus of 25-month-old mice that had undergone surgical gonadal removal, or sham surgery, and then were treated with 17aE2 or control diet. Mice underwent surgeries at 3 months of age and were treated with 17aE2 from 4 months to 25 months of age (Garratt *et al.* 2017).

Astrogliosis is correlated with increased expression of the glial fibrillary acidic protein (GFAP) (Thaler *et al.* 2012). In support of our previous report on 12-month-old mice (Sadagurski *et al.* 2017), 25-month-old sham males and females showed significantly different changes in hypothalamic astrogliosis with 17aE2 treatment, as can be seen by sex x drug interaction terms for GFAP^+^ astrocytes in the hypothalamus (p=0.02). Sham males responded to 17aE2 treatment with reduced numbers of GFAP^+^ astrocytes in the hypothalamus, with numbers comparable to sham or 17aE2 treated 25-months-old females (Fig 1). Comparing the response to 17aE2 in sham and castrated males, there was a significant interaction between surgery and drug treatment (p=0.004), demonstrating that castration diminished the response to 17aE2 in male mice. While for females the interaction between surgery and drug was not significant, ovariectomy significantly increased GFAP levels in both control and 17aE2 treated animals (p=0.004 for the main effect of surgery in a 2-way ANOVA including surgery type (sham or OVX) and treatment ((control or 17aE2) as independent factors) (Figure 1A and B).

**Figure 1:**
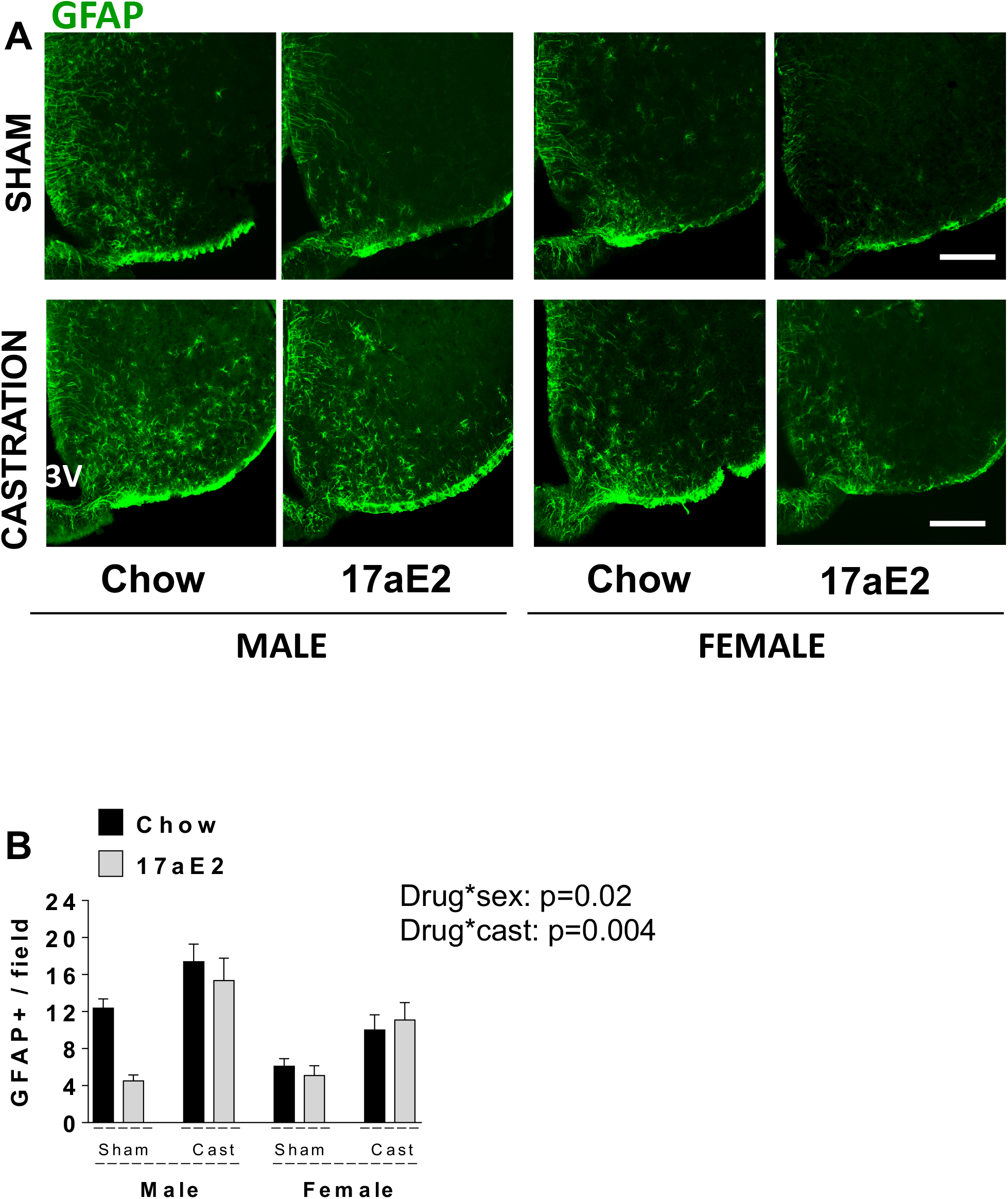
Hypothalamic astrogliosis in 17aE2 treated mice as measured by GFAP^+^ cells. Brain sections of 25-month-old male and female mice were analyzed for hypothalamic astrocytes. (A) Representative images showing immunostaining in the arcuate nucleus of hypothalamus (ARC) of chow-fed control and 17aE2 treated mice, castrated or ovariectomized. Scale bars: 200 μm, 3V, third ventricle. (B) Numbers of cells immunoreactive for GFAP in the ARC from indicated male and female mice; error bars show SEM for n = 5 mice of each type. The p-value represents the interaction term in a two-factor ANOVA, testing whether the drug effect differs between male and female mice (drug x sex: p=0.02), and its effect on castration (drug x cast: p=0.004).

Animal and human work demonstrate that changes in CA3 and dentate gyrus (DG) regions of the hippocampus are linked to age-related memory disorders (Cerbai *et al.* 2012; Lana *et al.* 2016). However, in our previous studies, we did not detect consistent effects of 17aE2 on astrogliosis in the hippocampus of the 12-month-old drug-treated mice (Sadagurski *et al.* 2017). In contrast, an assessment of GFAP^+^ astrocytes in CA3 of 25-month-old mice revealed that 17aE2 treatment protected males from age-associated astrogliosis, without a significant effect on females (sex x drug interaction p=0.016) (Figure 2B and D). Interestingly, the numbers of GFAP^+^ astrocytes in CA3 significantly increased in ovariectomized females on control chow and was significantly reduced in OVX females treated with 17aE2, with sham and OVX females showing a significantly different response to 17aE2 treatment (significant interaction term p=0.036). This indicates that female ovarian hormones influence the response to 17aE2 in this brain region. We found no statistical evidence that castration significantly influenced the response to 17aE2 in males since the treatment by surgery status interaction term was non-significant (p=0.125). In the DG region of the hippocampus, there was also significant sex by treatment interaction for sham-operated (intact) mice treated with 17aE2 (p=0.001), with males showing a decrease in GFAP^+^ astrocytes in response to 17aE2, while females showed no change (Figure 2B and D). In contrast to CA3, in the DG male castration eliminated the protective effect of 17aE2 in males (p=0.018 for the interaction between surgery and treatment within males). The numbers of the GFAP positive astrocytes in ovariectomized females were not different from sham surgery females, and ovariectomized females also did not respond to 17aE2 treatment. The sexually dimorphic responses in different domains of the hippocampus to gonadal hormones suggest that both male and female gonadally derived hormones can influence the severity of astrogliosis in old mice in a region-dependent manner, and both sets of hormones can influence responsiveness to 17aE2.

**Figure 2:**
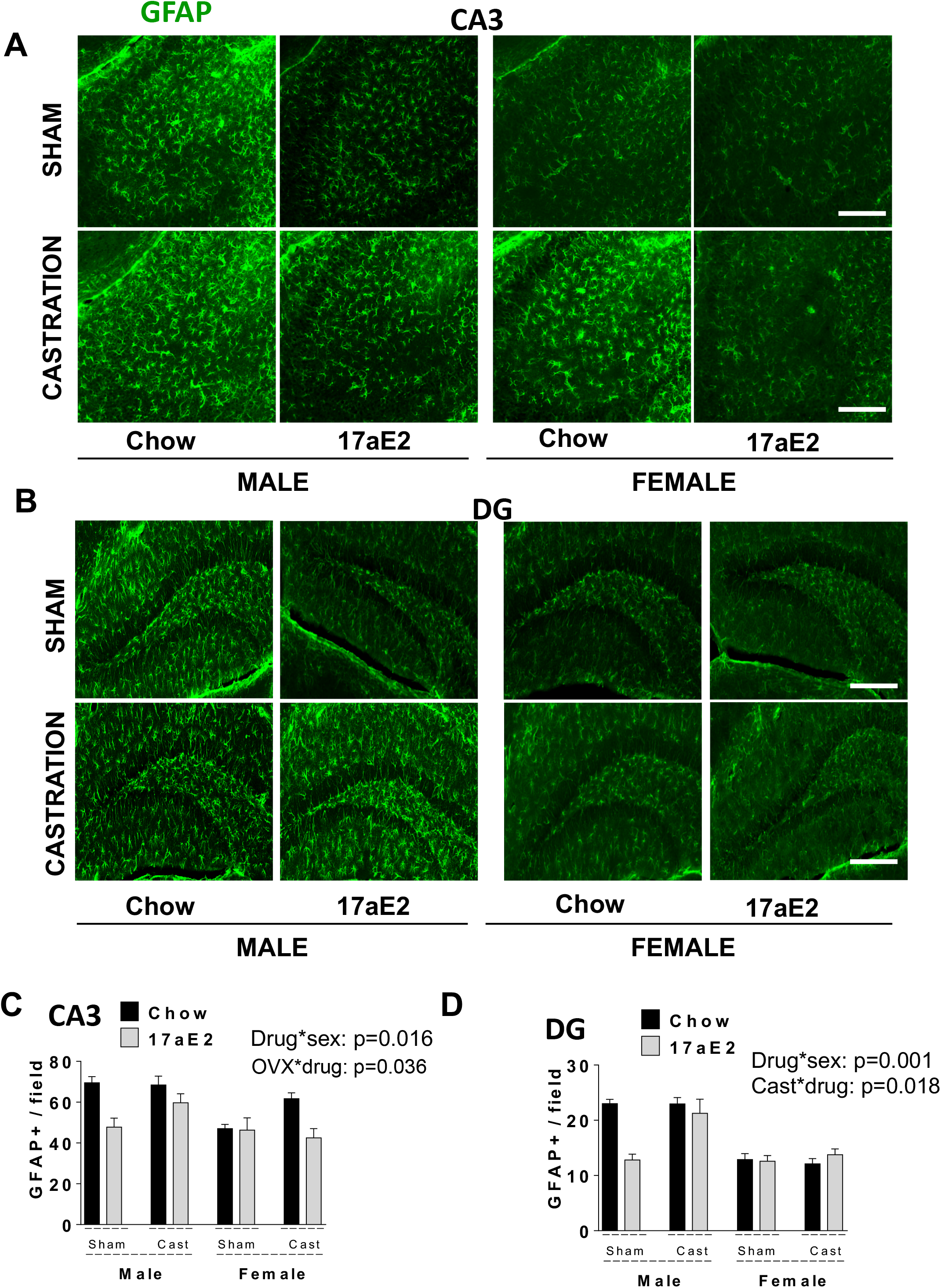
Astrogliosis in the hippocampus. Representative images of astrocytes identified by immunofluorescent detection of GFAP protein in coronal sections of (A) CA3 and (B) dentate gyrus (DG) area, respectively, obtained from 25-month-old male and female control and 17aE2 treated mice, castrated or ovariectomized. Scale bars: 200 μm, 3V, third ventricle. Quantification of GFAP staining represents the number of GFAP positive cells per field (error bars indicate SEM; n = 5 mice per group) in the (C) CA3 and (D) DG. The p-value represents the interaction term in a two-factor ANOVA, testing whether the drug effect differs between male and female mice (drug x sex: p=0.016 for CA3, p=0.001 for DG), and its effect on ovariectomy (drug x OVX: p=0.036 for CA3) and castration (drug x cast: p=0.018 for DG).

### Effect of sex hormones on microgliosis in 17aE2 treated mice

Using immunostaining for the microglia-specific ionized calcium-binding adaptor molecule 1 (Iba1), we next analyzed the number of microglial cells in the hypothalamus and the hippocampus (CA3 and DG). Consistent with the astrocyte data, the numbers of Iba1^+^ microglia were higher in the ARC of control males compared to females, and were reduced by 17aE2 treatment, with a significant interaction between sex and drug treatment compared to control (p=0.005) (Figure 3A and B). The effect of 17aE2 treatment was highly significant in males (p = 0.001) but was not significant in females. Looking at the effect of castration and ovariectomy on Iba1^+^ microglia and treatment responses within each sex, we observed no effect of castration on responses to 17aE2 or microglia levels within males (Figure 3A and B). However, the numbers of Iba1^+^ microglia were elevated in the ARC as a result of ovariectomy in both control and 17aE2 treated females, supporting the proposed role of estrogen on hypothalamic microglia inflammatory responses (Vegeto *et al.* 2003; Villa *et al.* 2016). There was no interaction between surgery status and drug treatment, with ovariectomized females also showing no change in Iba1^+^ microglia counts in the ARC with 17aE2 treatment. In the CA3 and DG, there was a similar sex-specific effect of the 17aE2 treatment on the astrogliosis of male but not female mice in intact (sham-operated) animals for the interaction effect of drug and sex (p=0.01) (Figure 4). The main effects of surgery for either sex were non-significant in CA3 and DG, indicating that castration and ovariectomy do not influence microgliosis in the hippocampus in a significant way in either sex (Figure 4A and B), although we note that the effect of ovariectomy was marginally non-significant (p=0.059). There was also no significant interaction between surgery and drug treatment within either sex, indicating that the manipulation of gonadal hormones as conducted in this study did not significantly influence 17aE2 responses in this brain area. Taken together our data show a sex-specific protective effect of 17aE2 on age-associated gliosis in the hypothalamus and the hippocampus, without a clear effect of gonadal removal on the 17aE2 responses in either sex.

**Figure 3:**
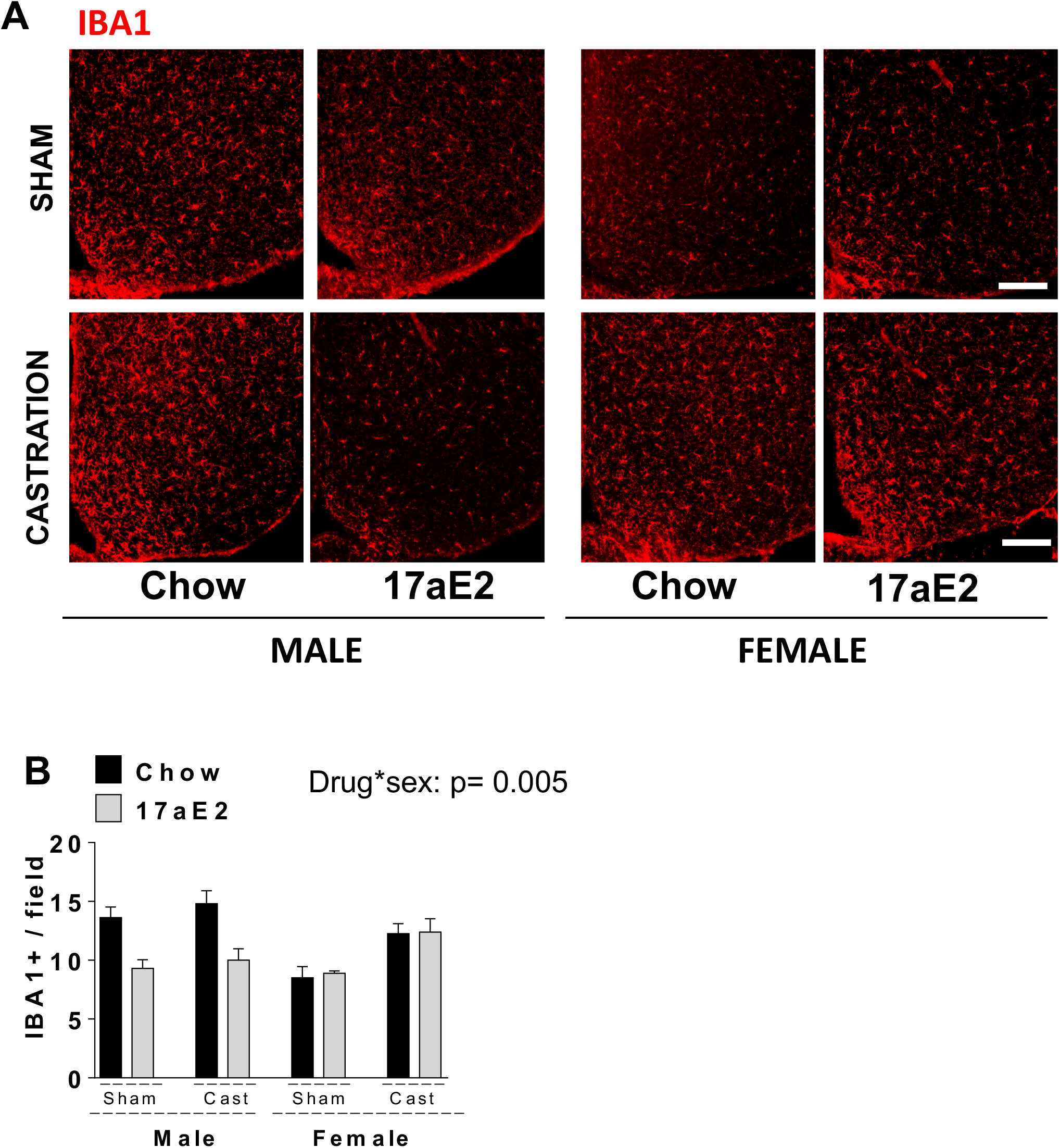
Hypothalamic microgliosis in 17aE2 treated mice as measured by Iba1^+^ cells. (A) Representative images showing immunostaining in the ARC of control and 17aE2 treated mice, castrated, or ovariectomized. Scale bars: 200 μm, 3V, third ventricle. (B) Numbers of cells immunoreactive for Iba1 in the ARC from indicated male and female mice; error bars show SEM for n = 5 mice of each type. The p-value represents the interaction term in a two-factor ANOVA, testing whether the drug effect differs between male and female mice (drug x sex: p=0.005), but no significant interaction between drug treatment and castration in either sex.

**Figure 4:**
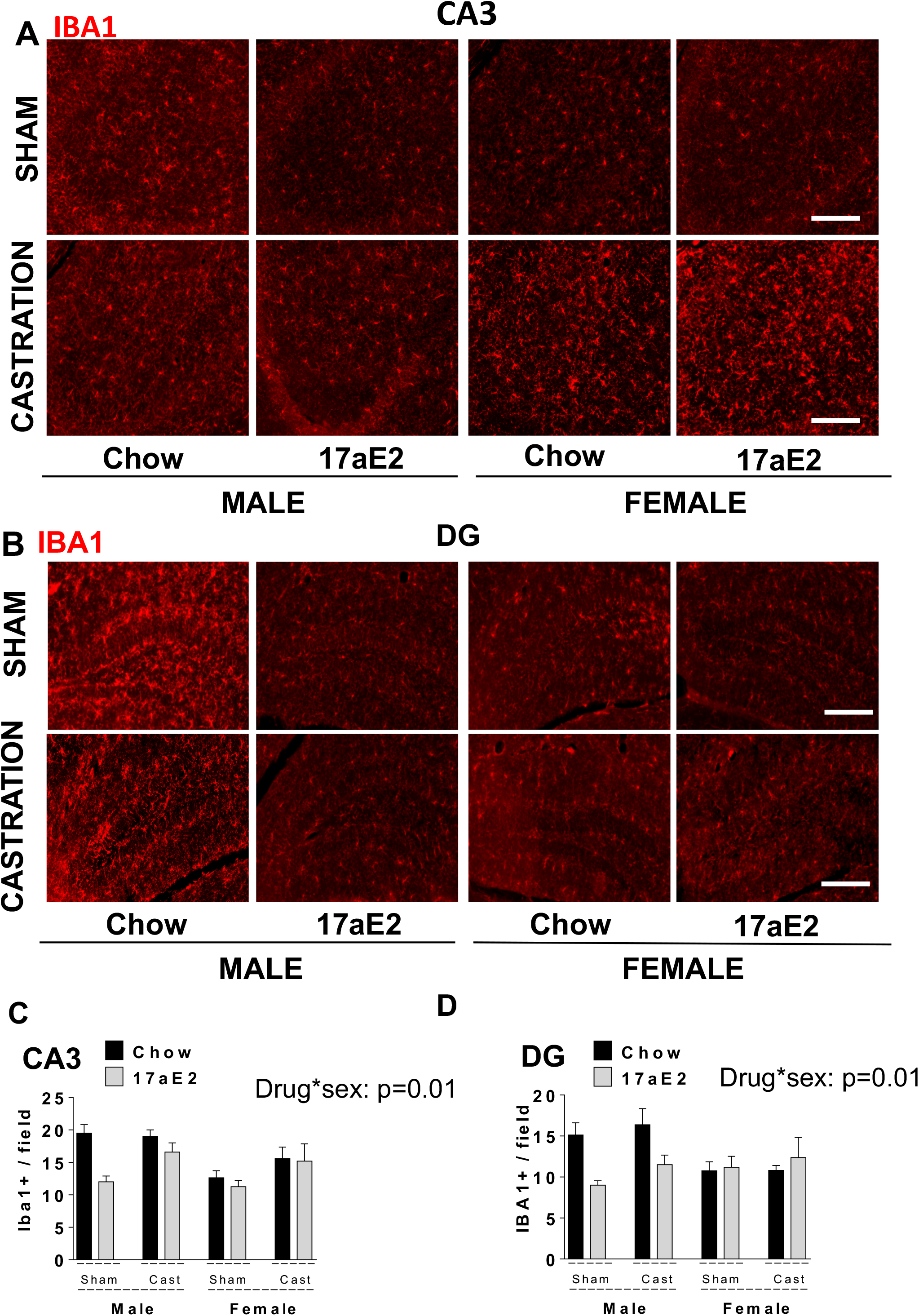
Microgliosis in the hippocampus. Representative images of microglia identified by immunofluorescent detection of Iba1 protein in coronal sections of (A) CA3 and (B) DG area, respectively, obtained from 25-month-old male and female control and 17aE2 treated mice, castrated or ovariectomized. Scale bars: 200 μm, 3V, third ventricle. Quantification of Iba1 staining represents the number of cells per field (error bars indicate SEM; n = 5 mice per group) in the (C) CA3 and (D) DG respectively. The p-value represents the interaction term in a two-factor ANOVA, testing whether the drug effect differs between male and female mice (drug x sex: p=0.01 for CA3, p=0.01 for DG), but no significant interaction between drug treatment and castration in either sex.

Female ovariectomy had an effect on astrogliosis and microgliosis in both regions, although the consistency of this effect was region-specific. These results suggest that gonadal hormone production from three months of age partly but not completely controls the sex-specific response to 17aE2 in these parameters.

### Effect of 17aE2 and sex hormones on hypothalamic ERα expression

In mice, ERα is a predominant steroid receptor expressed in the hippocampus and hypothalamus, and its expression levels are reduced during aging, particularly in the hippocampus (Foster 2012). ERα promotes neuronal survival and is more effective than ERβ in mediating ligand-independent transcription (Coleman *et al.* 2003). It was recently demonstrated that 17aE2 suppresses inflammation in adipocytes through its effects on the ERα and NFκB pathways (Santos *et al.* 2017). We hypothesized that the mechanism of action by which 17aE2 modulates neuroinflammatory responses in the hypothalamus might involve ERα activation. As can be seen in Figure 5, there is a male-specific increase in ERα expression in the ARC of 17aE2 treated males but not females. ERα expression in males was elevated to the levels similar to chow-fed or 17aE2 treated females and there was significant sex by treatment interaction in sham-surgery animals (p=0.001), indicating a different response in hypothalamic ERα expression to 17aE2 in males and females. For surgery effects within each sex, the surgery by treatment interaction was significant in male mice (p = 0.034), indicating that castrated males show a different response to 17aE2 than sham animals, with these animals showing no change in ERα abundance with 17aE2 treatment. In females, there was no interaction between surgery and 17aE2 treatment (p=0.295 for females), but the effect of surgery alone was significant (p = 0.001), indicating that females show a decrease in ERα expression in response to ovariectomy (Figure 5). We were not able to detect ERα in the hippocampus of old mice, in contrast to previous studies assessing hippocampal ERα signals in younger animals (Miller *et al.* 2005; Meyer & Korz 2013). This could suggest that ERα expression decreases in this area with age and that reduced inflammation in this brain region does not occur as a consequence of cell-autonomous ERα dependent processes, but involves additional mechanisms.

**Figure 5:**
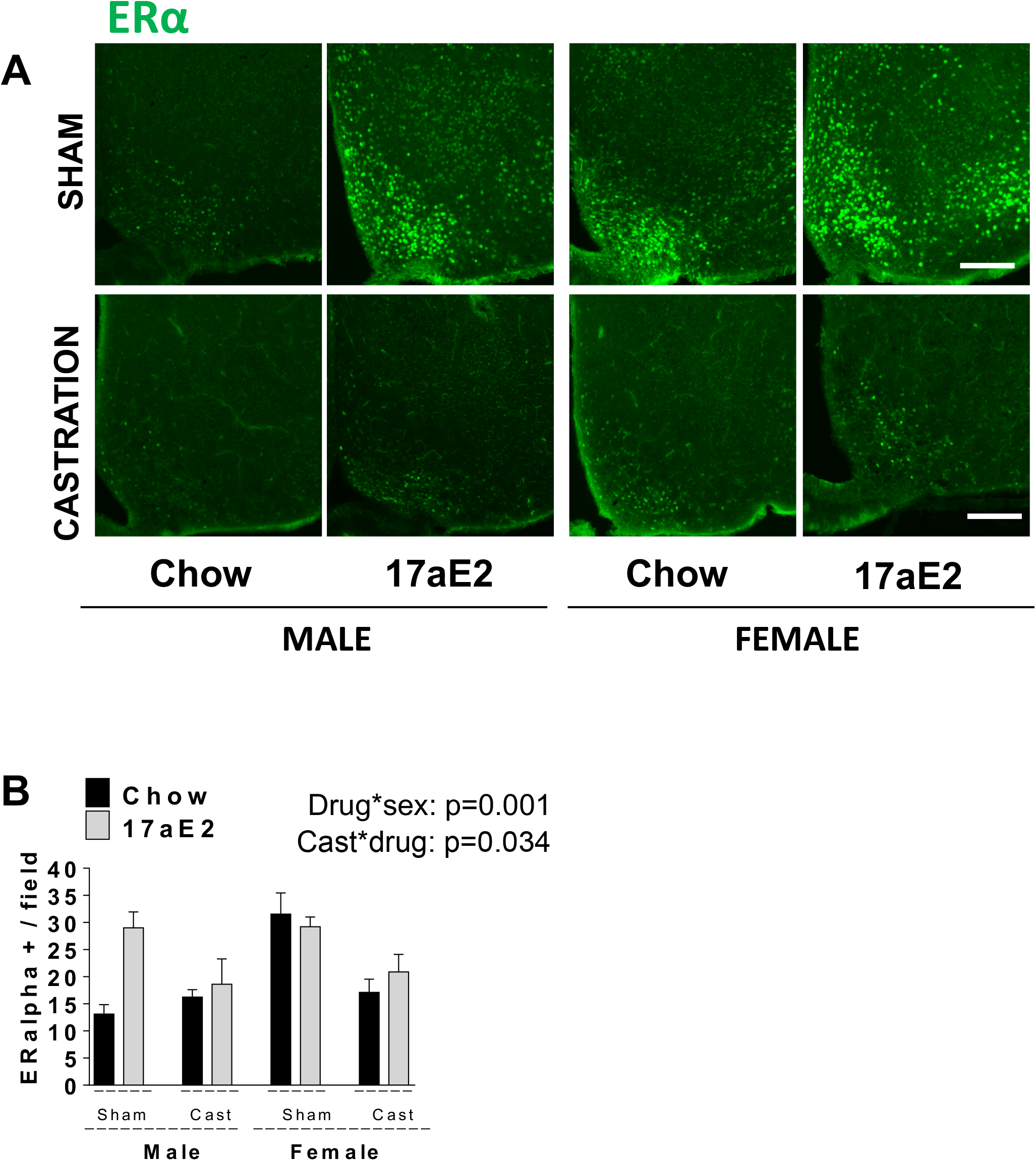
Hypothalamic ERα protein expression in 17aE2 treated mice. Brain sections of 25-month-old male and female mice were analyzed for hypothalamic ERα protein expression. (A) Representative images showing immunostaining in the ARC of control and 17aE2 treated mice castrated or ovariectomized. Scale bars: 200 μm, 3V, third ventricle. (B) Quantification of ERα protein in the ARC from indicated male and female mice; error bars show SEM for n = 5 mice of each type. The p-value represents the interaction term in a two-factor ANOVA, testing whether the drug effect differs between male and female mice (drug x sex: p=0.001), and its effect on castration (drug x cast: p=0.034).

### Effect of 17aE2 and sex hormones on hypothalamic mTOR signaling

mTOR signaling is involved in sex differences in aging and metabolism. Reduced mTORC1 signaling extends lifespan in female mice (Lamming *et al.* 2012), while reduced mTORC2 signaling reduces male lifespan without a significant effect on female lifespan (Lamming *et al.* 2014). Interestingly, 17aE2 treatment elevates hepatic mTORC2 activity in males, without a significant effect on mTOR1 signaling in either male or female mice (Garratt *et al.* 2017). In the ARC, mTORC1 signaling can mediate estrogen’s anorectic effects on energy homeostasis (Hu *et al.* 2016). We assessed S6 phosphorylation, one of the downstream substrates of mTORC1 in the ARC of 25-month-old mice. Consistent with the liver data (Garratt *et al.* 2017), S6 phosphorylation in the ARC did not significantly change with 17aE2 treatment, and there was no significant effect of sex (Figure 6). On the other hand, castration overall increased phosphorylation levels of pS6 in both chow and 17aE2 treated males, although there were surgery and sex by drug interaction (p=0.04; Figure 6), because 17aE2 treatment reduced pS6 specifically in castrated male mice (p=0.001). There was no overall effect of ovariectomy on pS6 and no interaction between ovariectomy and drug treatment within females.

**Figure 6:**
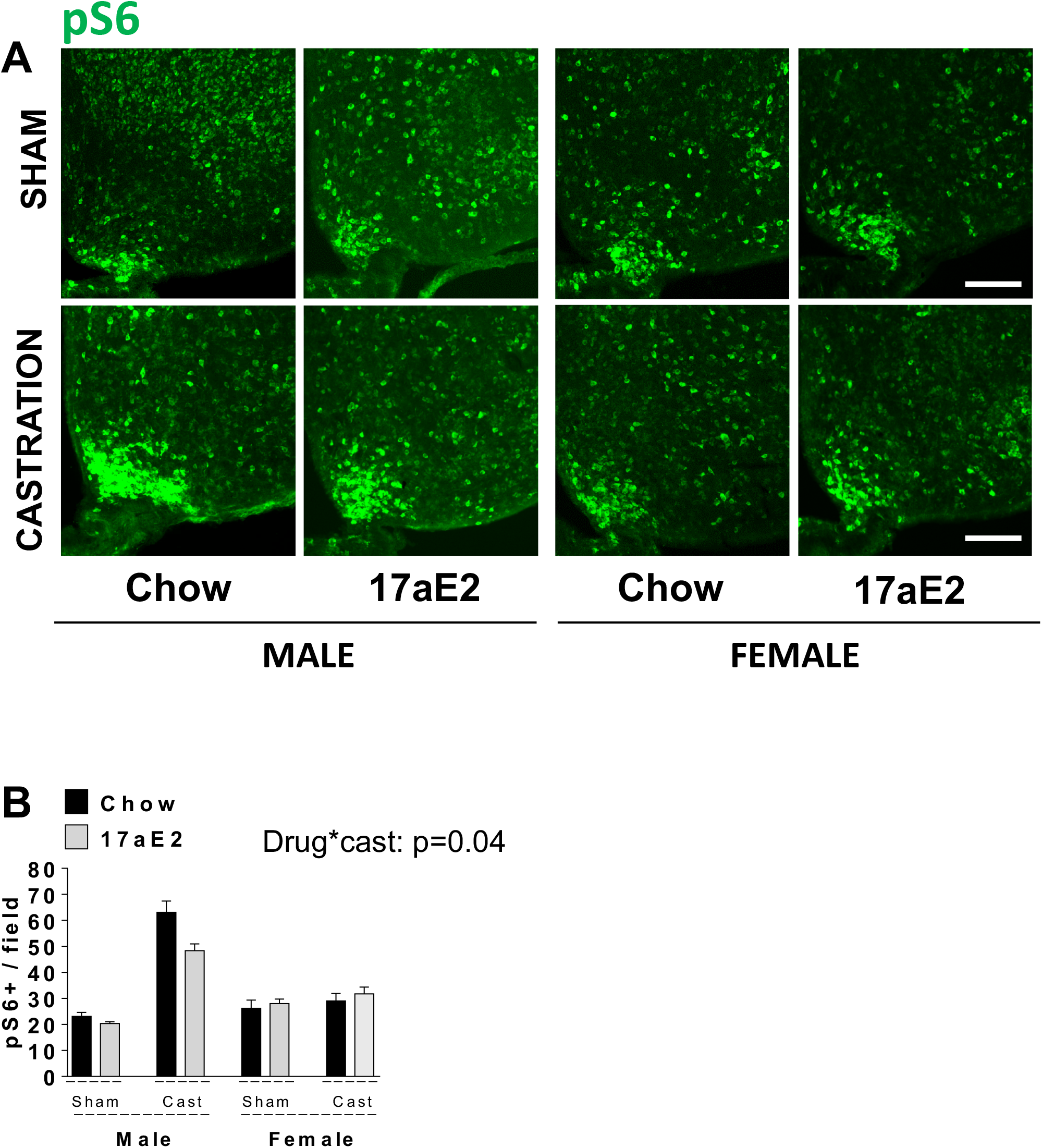
Hypothalamic pS6 protein expression in 17aE2 treated mice. Brain sections of 25-month-old male and female mice were analyzed for hypothalamic pS6 protein expression. (A) Representative images showing immunostaining in the ARC of control and 17aE2 treated mice. Scale bars: 200 μm, 3V, third ventricle. (B) Quantification of pS6 in the ARC from indicated male and female mice; error bars show SEM for n = 5 mice of each type. The p-value represents the interaction term in a two-factor ANOVA, between drug treatment and castration (drug x cast: p=0.04).

## Discussion

Our results show that sex-specific effects of 17aE2 on lifespan are reflected in reduced age-associated neuroinflammation specifically in males, within brain regions that are sensitive to neurodegeneration and metabolic imbalance. These sexually dimorphic neuroinflammatory responses are brain region-specific and are partially, but not entirely, dependent on gonadal hormone production. We provide evidence that the male-specific effects of 17aE2 are correlated with male-specific changes in ERα expression in the ARC, responses that are not observed in females, or castrated males.

Castrated males do not show the decline in age-associated gliosis produced by 17aE2-treatment in intact males (Fig. 1), and do not respond to 17aE2 with an increase in ERα expression in the ARC, as seen in intact males (Fig. 5). Male gonads, probably via testosterone production, therefore contribute to the sexually dimorphic responses elicited by 17aE2, with castrated males showing the lack of drug effect that is observed in sham females. Ovariectomy activates age-associated hypothalamic gliosis in females that are otherwise protected from hypothalamic inflammation. A similar increase in age-associated gliosis is observed in CA3 after ovariectomy, but this hormonal effect is not apparent in DG. Each of these areas of the hippocampus has a distinctive function that is differentially affected by the brain aging process (Dillon *et al.* 2017). Interestingly, in the CA3, 17aE2 also reduced gliosis in OVX females, indicating a role of both male and female sex-hormones in the sex-specific responses to 17aE2, depending on the brain region. Neuroinflammation has been reported to take place prior to overt neuron loss in various animal models of age-related neurodegeneration (Tabuchi *et al.* 2009), raising the possibility that the early use of drugs that can delay neuroinflammation could play a critical role in mitigating disease progression. In support, it has been suggested that 17aE2 could be effective in the protection against oxidative stress, amyloid toxicity, and Parkinson’s and Alzheimer’s diseases (Dykens et al. 2005).

We could not detect the expression of ERα in the hippocampus of old brains of either sex. Although no report has specifically examined the effect of age on the expression of ERα in the hippocampus of mice, there was 80% decline in ERα expression in the hippocampus of aged rats (Mehra *et al.* 2005), which may explain why ERα was below the threshold for detection in this brain region in our study. Since knockout of ERα expression in mice is associated with a decrease in the expression of genes critical for hippocampal function during aging (Han *et al.* 2013), the loss of detectable ERα in these brain regions during aging could contribute to impairing brain function stemming from this region. Indeed, estrogen’s effects on cognition are reduced with aging, and it has been suggested that this could be caused by altered expression of ERα in the hippocampus (Bohacek & Daniel 2009).

Our previous study did not detect a significant effect of 17αE2 on neuroinflammatory responses in the hippocampus of 12-month-old mice (Sadagurski *et al.* 2017). However, we now show that by 25-months of age, both astrogliosis and microgliosis are significantly reduced by 17αE2 in CA3 and DG in males, though not in females. Since we do not detect ERα in these regions at this time point, our results suggest that either other hormone receptors/signaling processes are involved in causing changes in this brain area, or that ERα mediates the signaling effects of 17αE2, but effects in the hippocampus are indirect and occur as a consequence of ERα activation occurring in a different brain region.

ERα is strongly expressed in the ARC and plays an important role in metabolic regulation. Female but not male mice with ERα selectively deleted in Pro-opiomelanocortin (POMC) neurons in the ARC are hyperphagic and develop modest obesity (Xu *et al.* 2011). Our data shows that in response to 17aE2 treatment, ERα expression levels are elevated in the ARC of male mice and become similar to the ERα levels in chow-fed or 17aE2 treated females. ERα is expressed in astrocytes and can activate several neuroprotective mechanisms including the control of neuroinflammation (Vegeto *et al.* 2003). These data could potentially imply that the ability of 17aE2 to decrease neuroinflammatory responses in the hypothalamus depends on interactions with ERα, a hypothesis worthy of future investigation. In support, castration reduced ERα levels in the ARC, while numbers of activated astrocytes increased, and 17aE2 did not reduce astrocytes in castrated males. Similarly, 17aE2 suppressed inflammation in primary adipocytes through their effects on ERα in a sex-specific manner with males being more sensitive to its effects (Santos *et al.* 2017).

A recent study suggested that the central actions of estrogen on energy balance are at least partially mediated by the selective modulation of mTOR pathway through ERα in the ARC (Gonzalez-Garcia *et al.* 2018). We show that castration reduced the expression levels of hypothalamic ERα in 17aE2 treated males, and significantly increased mTORC1 signaling in the ARC. While the molecular mechanisms underlying this relationship require further investigation, such an interaction could be related to the altered activity of mTORC2 signaling in the ARC neurons (Chellappa *et al.* 2019). Understanding the effect of hypothalamic mTOR signaling in the control of neuroinflammatory responses may provide significant insight into the causes of sexspecific differences in aging.

Our previous work has shown that a variety of other metabolic and anti-aging responses to 17aE2 occur in males but not females, and that these sex-specific responses are not seen in castrated males (Garratt *et al.* 2017). Improvements in glucose tolerance and enhanced hepatic mTORC2 signaling with 17aE2 treatment are observed in male mice only, and these effects are inhibited in castrated males. Furthermore, 17aE2 treatment improves male grip strength and capacity to balance on a rotarod, and these improvements are not seen in castrated male mice treated with 17aE2 (Garratt *et al.* 2019). The hypothalamus is a major central regulator of peripheral metabolism, and the regulation of metabolism via this region is influenced by inflammatory responses (Zhang *et al.* 2013). Thus, it is possible that reduced hypothalamic inflammation by 17aE2 could promote the metabolic and health benefits of this drug observed in other regions of the body.

It has previously been shown that some responses to 17aE2, particularly changes in feeding behavior, are dependent on the presence of functional POMC neurons (Steyn *et al.* 2018), although other metabolic responses to 17aE2 remain intact. The ARC of the hypothalamus, where we observe changes in the expression of ERα and reduced inflammation with 17aE2, contains POMC neurons. This could suggest a connection between activation of these neurons via ERα, altered feeding behavior, and also changes in the abundance of astrocytes and microglial in close proximity. The ARC also contains additional neuronal populations involved in metabolic homeostasis, many of which express ERα (Olofsson *et al.* 2009; Herber *et al.* 2019), which are also reasonable candidates through which 17aE2 may elicit its metabolic effects. Ultimately, hypothalamic processes are strongly implicated in the control of systemic aging, particularly in response to inflammation (Zhang *et al.* 2013). Therefore careful dissection of the central signaling pathways and neuronal populations through which 17aE2 influences metabolic homeostasis, and ultimately lifespan, could provide a key insight into lifespan control. The alternative hypothesis is that 17aE2 acts at peripheral sites like the liver, muscle, or adipose tissue, potentially helping to maintain metabolic regulation and glucose control, and this reduces hypothalamic inflammation. Distinguishing the central and peripheral signaling targets of 17aE2, and the receptors involved, would be crucial in the development of targeted androgen and estrogen pathway modulators that help to slow aging while minimizing adverse health outcomes.

## Methods

### Animals

Procedures involved in this study were approved by the University of Michigan Committee on the Use and Care of Animals (UCUCA). UM-HET3 mice were produced as previously described in detail (Miller *et al.* 2007). For breeding cages, we used Purina 5008 mouse chow. After weaning all animals were fed Purina 5LG6 until 4 months of age. Animals were maintained under temperature- and light-controlled conditions (20-23°C, 12-h light-dark cycle).

### Surgical procedures

At three months of age, all animals went through castration, ovariectomy, or a sham procedure. All animals were anesthetized by injection of 250mg/kg tribromoethanol, and given a single pre-operative injection of the analgesic carprofen, at 5 mg/kg.

### Castration and sham castration

After surgical preparation, an incision was made in the caudal end of each scrotal sac, the testicle was pulled through the incision by gentle traction, and the blood vessels, vas deferens, and deferential vessels were clamped and sutured. The incision was closed with tissue adhesive. For sham surgery, the testicles were exteriorized and then replaced in the scrotum, without being ligated or excised.

### Ovariectomy or sham ovariectomy

After surgical preparation, an incision was made on the left side perpendicular to the vertebral column approximately midway between the iliac crest and the last rib. The ovarian fat pad was grasped and exteriorized. The pedicle under the ovarian blood vessels and fat pad under the ovary were grasped and crushed, the pedicle cut on the ovary side and the ovary removed, and the blood vessels tied with absorbable suture. The abdominal wall was closed with absorbable suture and skin was closed with staples. The procedure was then repeated on the opposite side. For sham ovariectomy, animals underwent the same surgical procedure, but the ovary and fat pad were exteriorized and replaced without being excised.

### Experimental diets

17aE2 was purchased from Steraloids Inc. (Newport, RI, USA) and mixed at a dose of 14.4 milligrams per kilogram diet (14.4 ppm). Animals from each surgical type were randomly allocated to control or 17aE2 treatment. Control animals were maintained on 5LG6 while the treatment group were fed the 17aE2 diet continuously from 4 months of age.

### Perfusion and immunolabeling

Mice were anesthetized (IP) with Avertin and transcardially perfused with phosphate-buffered saline (PBS) (pH 7.5) followed by 4% paraformaldehyde (PFA). Brains were post-fixed, dehydrated, and then sectioned coronally (30 μm) using a sliding microtome, followed by immunofluorescent analysis as previously described (Sadagurski *et al.* 2017). Brain sections were washed with PBS several times; blocked for 2h in 0.3% Triton X-100 with 3% normal donkey serum in PBS; and then stained with the following primary antibodies overnight: rabbit anti-GFAP (1:1000; Millipore,Temecula, CA) and rabbit anti-Iba1 (1:1000; Wako, Richmond, VA). For rabbit anti ER alpha (1:1000; Santa Cruz Biotechnology, CA) and rabbit anti pS6 (1:100 Cell signaling, MA) immunostaining, sections were pretreated for 20 min in 0.5% NaOH and 0.5% H_2_O_2_. in PBS, followed by immersion in 0.3% glycine for 10 min. Sections were then placed in 0.03% SDS for 10 min and placed in 4% normal serum plus 0.4% Triton X-100 plus 1% BSA for 20 min and then stained with primary antibodies overnight. All floating brain sections were washed with PBS several times; and incubated with AlexaFluor-conjugated secondary antibodies for 2h (Invitrogen, Carlsbad, CA). Sections were mounted onto Superfrost Plus slides (Fisher Scientific, Hudson, NH) and cover slips added with ProLong Antifade mounting medium (Invitrogen, Carlsbad, CA). Microscopic images were obtained using an Olympus FluoView 500 Laser Scanning Confocal Microscope (Olympus, Center Valley, PA) equipped with a 10x, 20x and 40x objectives.

### Quantification

For quantification of immunoreactive cells, images of matched brain areas were taken from at least three sections containing the hypothalamus for each brain between bregma −0.82 mm to −2.4 mm (according to the Franklin mouse brain atlas). Brain slices were taken at the same distance from bregma. Serial brain sections across the hypothalamus were made at 30 μm thickness, and every five sections were represented by one section with staining and cell counting. All sections were arranged from rostral to caudal to examine the distribution of labeled glial cells. Iba1 and GFAP positive cells and ER alpha and pS6 protein staining were counted using ImageJ software with DAPI (nuclear stain). The average of the total number of cells/field of view was used for statistical analysis as described previously (Sadagurski *et al.* 2017).

### Statistical analysis

For each measured parameter, we conducted a two factor ANOVA, using the general linear model function and a full factorial model, which included an effect of treatment (comparing control to either ACA or treatment), an effect of sex (male or female) and an interaction between sex and treatment. When testing for the effect of gonadectomy on treatment responses within each sex, we included an effect of treatment, an effect of surgery (gonadectomised or not), and an interaction between surgery and treatment. Data were transformed where necessary to conform to assumptions of normality. IBM SPSS v.21 was used for statistical analysis.

## Abbreviations

ACA: acarbose
17aE2: 17-a-estradiol
CNS: central nervous system
ERα: estrogen receptor alpha
ARC: arcuate nucleus of the hypothalamus

## Acknowledgments

We thank Amanda Keedle, Lynn Winkelman, Sabrina Van Roekel, Roxann Alonso, Gillian Cady, and Natalie Perry for husbandry and technical assistance.

